# Spray Dried Inulin–Montmorillonite Hybrids Alleviate High-Fat Diet–Induced Inflammatory and Metabolic Dysregulation in Rats

**DOI:** 10.64898/2026.01.22.701174

**Authors:** Amin Ariaee, Alex Hunter, Anthony Wignall, Kristen Bremmell, Clive Prestidge, Paul Joyce

## Abstract

Obesity-related metabolic disorders are linked to excessive dietary lipid absorption and gut microbiota imbalances, particularly under high-fat diet (HFD) conditions. This study evaluates a spray-dried hybrid of inulin and montmorillonite (INU-MMT) designed to concurrently restrict intestinal lipid digestion and modulate the gut microbiota. Using an *in vitro* simulated intestinal lipolysis model, INU-MMT significantly reduced free fatty acid (FFA) release from medium-chain triglycerides by 4.0-fold compared to HFD conditions, outperforming INU and MMT individually. This superior inhibition is attributed to INU’s ability to prevent MMT aggregation, resulting in smaller, more dispersed particles with enhanced lipid-binding capacity. In a 21-day *in vivo* study in HFD-fed rats, INU-MMT (1g/kg bodyweight/d) supplementation significantly attenuated cumulative weight gain by 4.7% compared to the HFD control, exceeding the effects of INU (2.0%) and MMT (1.5%) alone. 16S rRNA gene sequencing of fecal samples revealed improved gut microbial diversity (Simpson’s index, p = 0.0161) and enrichment of health-associated taxa including *Peptostreptococcaceae* (8-fold), *Ruminococcaceae* (3.5-fold), *Akkermansiaceae* (2.5-fold), and *Eggerthellaceae* (7.7-fold). Beta diversity analysis highlighted that INU-MMT induced a distinct microbial composition from HFD and INU groups (PERMANOVA, adjusted p < 0.05), driven largely by MMT. Predictive metagenomic analysis using the Phylogenetic Investigation of Communities by Reconstruction of Unobserved States 2 (PICRUSt2) software demonstrated a 98% reduction in microbial triacylglycerol lipase abundance, aligning with the observed *in vitro* lipolysis suppression results. These findings highlight the dual-mechanistic potential of INU-MMT in managing diet-induced obesity by targeting lipid digestion and imbalances within the gut microbiota.

**Graphical abstract:** 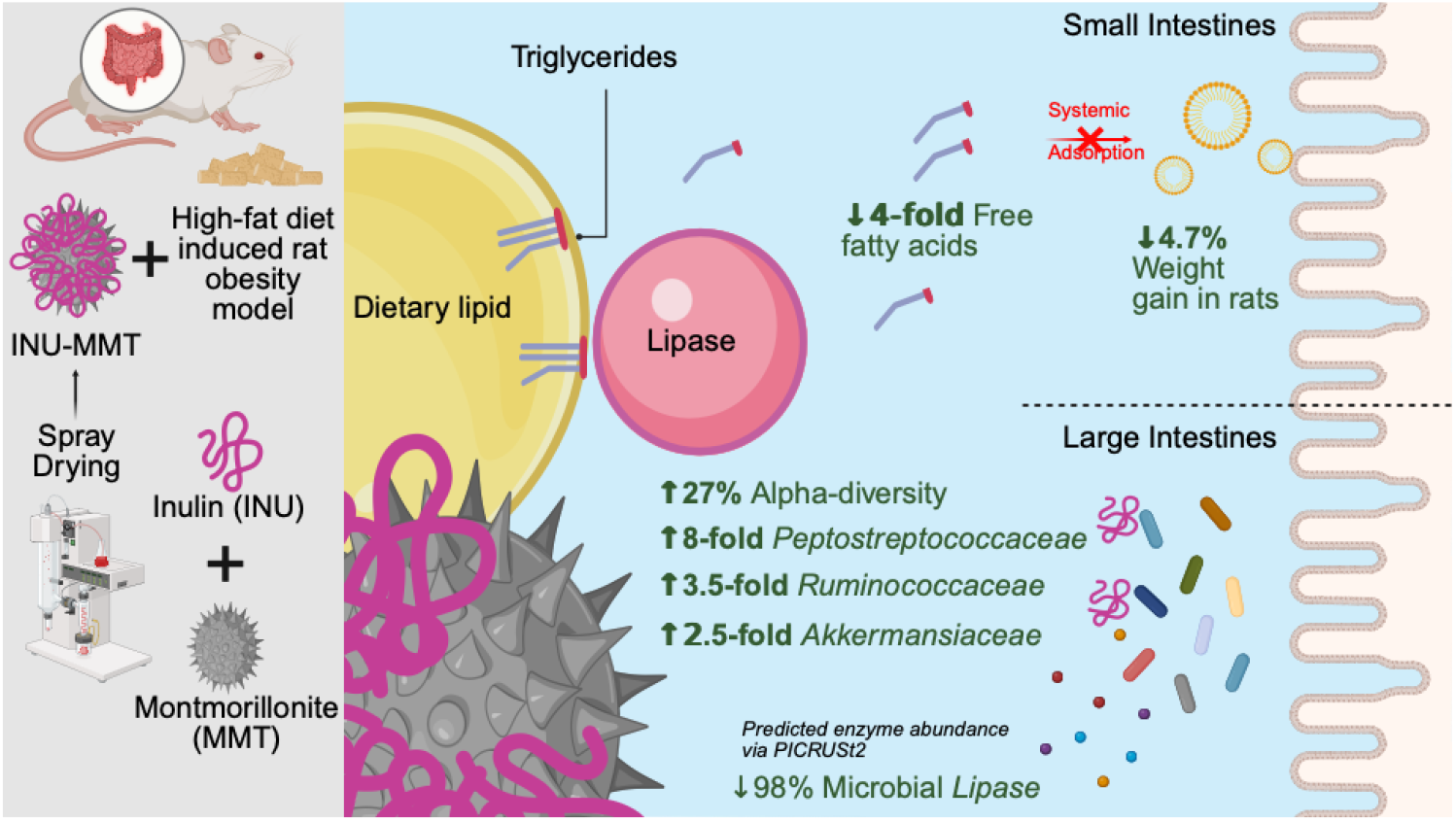

**Highlights:**

- Spray-dried INU-MMT restricts FFA release 4-fold in intestinal conditions
- Hybrid outperforms INU and MMT in reducing HFD-induced weight gain in rats
- Promotes beneficial microbiota shifts and key SCFA-producing taxa
- Suppressed predicted microbial lipase levels by 98% with INU-MMT treatment
- INU-MMT offers a multi-mechanistic strategy for future metabolic disease therapies.

## 1. Introduction

The global rise in obesity has become a major public health challenge, closely associated with metabolic disorders such as type 2 diabetes, cardiovascular disease, and non-alcoholic fatty liver disease [1, 2]. Among the various mechanisms driving metabolic dysregulation, excessive dietary fat intake and pathogenic shifts in the gut microbiota have emerged as key, yet underexplored therapeutic targets for metabolic dysregulation [3, 4]. These disruptions, commonly associated with high-fat diets (HFD) in developed countries, contribute to systemic inflammation and metabolic imbalance [5, 6].

Growing evidence supports the potential of functional biomaterials to restore gut microbiota balance and improve metabolic outcomes in diet-induced obesity (DIO) models [7, 8]. Among these, inulin (INU), a fermentable prebiotic fiber, is selectively metabolized by beneficial gut microbes expressing enzymes such as exo-inulinase and β-fructosidase [9]. This fermentation process generates short-chain fatty acids (SCFAs), which enhance gut barrier integrity, reduce inflammation, and stimulate the secretion of satiety hormones such as glucagon-like peptide-1 (GLP-1) and peptide YY (PYY) [10-17]. However, the metabolic benefits of INU are often modest under HFD conditions, which may in part be due to dietary shifts in microbial composition reducing the abundance of INU-fermenting taxa, thereby diminishing SCFA production and downstream effects [7, 18, 19]. Therefore, combining INU with complementary gut-active materials may help overcome these limitations and enhance therapeutic efficacy.

One biomaterial with demonstrated efficacy in reducing the metabolic burden of HFDs is montmorillonite (MMT). MMT is aluminosilicate clay with strong binding capacities for pollutants and toxins and has more recently been explored for its binding of dietary lipids and their lipolytic products [20-23]. These properties have been correlated with MMT administration reducing weight gain in diet-induced animal models of obesity [24-26]. Moreover, spray dried MMT microparticles have been demonstrated to adsorb significant proportions of lipolytic products and perform comparably to orlistat in inhibiting weight gain in rodents [22, 26]. The current study further explores the potential of combining INU and MMT into a spray-dried hybrid (INU-MMT) particulate material, aiming to leverage the prebiotic and anti-inflammatory properties of INU with the lipid-binding and microbial regulatory functions of MMT. We hypothesize that the INU-MMT hybrid will provide synergistic effects in mitigating diet-induced metabolic dysfunction by simultaneously targeting the restriction of lipid digestion pathways and inducing shifts in the gut microbiota towards metabolic health.

## Methods and Materials

### 1.1 Materials

INU with a degree of polymerization 14 (determined by ^1^ H NMR, data not presented) and MMT were sourced from Pharmako Biotechnologies (Australia). Medium-chain triglycerides (MCT; Miglyol 812) were obtained from Hamilton Laboratories (Australia), while fed-state simulated intestinal fluid (FeSSIF) powder was purchased from Biorelevant.com Ltd (United Kingdom). *Lipase* from *Candida antarctica* (6000 TBU/mL), starch, cellulose, sodium hydroxide, glacial acetic acid, sodium chloride, and phosphate buffered saline (PBS) tablets were supplied by Sigma-Aldrich (Australia). Porcine pancreatin extract (8× USP) was obtained from MP Biomedicals (Australia). Four-week-old male Sprague-Dawley rats were sourced from the Animal Resources Centre (Australia). All chemicals and solvents were analytical grade, and Milli-Q water was used throughout.

### 1.2 Preparation of INU-MMT hybrids using spray drying

INU-MMT hybrids (at a 1:1 mass ratio) were prepared via spray drying, following established methods [27, 28]. Briefly, a 2% w/v INU dispersion was made by dissolving 20 g of INU in 1 L of Milli-Q water and stirring for 30 minutes at room temperature. Simultaneously, 20 g of MMT was dispersed in 1 L of Milli-Q water under the same conditions. The aqueous dispersions were combined in a 1:1 mass ratio and spray dried using a Büchi B-290 Mini Spray-Dryer (Germany) with an inlet temperature of 200°C, outlet temperature of 105°C, aspirator at 100%, nozzle cleaning at 9, compressed air flow at 40 mm, and product flow at 7 mL/min.

### 1.3 Surface morphology examination via scanning electron microscopy

The size and morphology of INU-MMT hybrids and their precursor materials were analyzed using a Zeiss Crossbeam 540 scanning electron microscope (SEM) (Germany). Samples were mounted on aluminum stubs using carbon tape and coated with a 10–20 nm platinum layer via sputter coating. Imaging was conducted at an accelerating voltage of 1–2 kV, and micrographs were processed using AztecOne software (Oxford Instruments, United Kingdom). For cross-sectional SEM analysis, focused ion beam (FIB) milling was performed using a Helios NanoLab FIB-SEM instrument (Russia). Spray-dried INU-MMT particles were sectioned using a gallium ion beam under vacuum, and the exposed internal structures were imaged under high-resolution SEM mode to visualize internal morphology.

### 1.4 Determination of particle size using laser diffraction

Particle size analysis of INU, MMT, and spray-dried INU-MMT hybrids was conducted using Mastersizer laser diffraction equipment from Malvern Instruments (United Kingdom). Samples were dispersed in an aqueous buffer (pH 6.5) under continuous stirring for 60 minutes to simulate small intestinal conditions. The particle size distribution was assessed using the D50 value, representing the median particle diameter. Measurements were performed in triplicate. INU-MMT’s adsorption of dietary lipids via confocal laser scanning microscopy

Confocal fluorescence imaging was performed using a Zeiss Elyra PS-1 Microscope (Germany) to visualize *in vitro* lipolysis samples. Digestible dietary lipid, being medium-chain triglycerides (MCT), and INU-MMT were stained with coumarin 6 and rhodamine B, respectively. After 60 minutes of lipolysis, sample aliquots were collected to examine the adsorption behavior of the spray-dried clay particles. Confocal images were captured at an emission wavelength of 497 nm for coumarin 6 (excitation 488 nm) and 588 nm for rhodamine B (excitation 514 nm), resulting in green and red fluorescence, respectively.

### 1.5. Simulated intestinal lipid digestion assay

An *in vitro* intestinal lipolysis model was used to assess the effects of INU, MMT, and spray-dried INU-MMT hybrids on lipid digestion under fed-state intestinal conditions (pH 6.5) [23, 26]. Simulated small intestinal (pH 6.5) media were prepared using a 50 mM Tris–maleate buffer and used within 48 hours. Pancreatin extract was prepared by dissolving 2 g of pancreatin powder in 10 mL of FeSSIF (pH 6.5), followed by centrifugation at 2268 × g for 20 minutes at 4 °C. The supernatant was collected and stored at 4 °C until use. MCT (625 mg) was dispersed in 20 mL of FeSSIF by stirring at 600 rpm for 10 minutes at 37 °C. INU, MMT, or INU-MMT hybrids were added at 10% w/w relative to lipid content.

The pH was maintained at 6.5 ± 0.01 using 0.1 M sodium hydroxide or hydrochloric acid. Lipolysis was initiated by adding 2 mL of pancreatin extract (2000 tributyrin units), and free fatty acid (FFA) release was monitored for 25 minutes using the 902 Titrando pH-stat titrator from Metrohm (Switzerland), which maintained a constant pH of 6.5 by titrating 0.6 M sodium hydroxide.

### 1.6. *In vivo* study design

The *in vivo* study was approved by the Animal Ethics Committee at the University of South Australia (approval #U24-21) and conducted in accordance with the NIH Principles of Laboratory Animal Care (NIH publication #85-23, revised 1985), the Australian Code for the Care and Use of Animals for Scientific Purposes (8th edition, 2013, revised 2021), and ARRIVE 2.0 guidelines for *in vivo* research [29].

Four-week-old male Sprague-Dawley rats were housed in pairs under controlled conditions (12-hour light/dark cycle, temperature, humidity, and pressure) and acclimated for one week before the study. They were then placed on a HFD (22% fat/44% energy from fat) and assigned to treatment groups, as conducted in previous studies [7, 8]. Each treatment was dispersed in PBS and administered daily via oral gavage between 16:00 and 18:00 over a three-week period, aligning with rodents’ natural nocturnal feeding behavior. The dosage treated was 1g/kg of bodyweight per day. Body weight was recorded daily before dosing.

At the end of the treatment phase, rats were anesthetized with isoflurane before euthanasia by cervical dislocation.

### 1.7. 16s rRNA gene sequencing of fecal samples

After humane euthanasia, fecal samples were collected and analyzed at the Australian Genomics Research Facility (Brisbane, Australia) for DNA extraction and 16S rRNA sequencing, focusing on the V3–V4 hypervariable regions. Sequence clustering into operational taxonomic units (OTUs) was performed at a 97% similarity threshold using QIIME 2, with taxonomic classification based on the Silva reference database (Release 138.1). The QIAGEN CLC Genomics Workbench (Version 23.0.4) and QMI-PTDB TaxPro index (June 2021) were used for further taxonomic assignments.

Microbial diversity was assessed through alpha diversity metrics, including Shannon, Chao1, and Simpson indices, along with total OTU counts. Beta diversity was evaluated using Bray-Curtis dissimilarity, Jaccard indices, and Euclidean distances. Statistical significance in beta diversity was determined using Permutational Multivariate ANOVA (PERMANOVA), with Principal Coordinates Analysis (PCoA) plots displaying 95% confidence ellipses to illustrate group clustering and variability. Samples that did not meet quality control standards set by the Qiagen Metagenomic module were excluded, resulting in a final sample size of n=5 per group.

### 1.8 16s rRNA metagenomic predictions using PICRUSt2

PICRUSt2 (Phylogenetic Investigation of Communities by Reconstruction of Unobserved States) was utilized as a QIIME2 plugin to predict enzyme presence and metabolic pathway abundances in each sample [30]. Enzyme predictions were generated using the Enzyme Commission (EC) database (Nov 2023).

### 1.9 Statistical analysis

All experimental data, excluding 16S rRNA gene sequencing results, were analyzed using GraphPad Prism Version 10.2.0 (Boston, MA, USA). Before performing statistical analyses, data and residuals were tested for normality using the Shapiro-Wilk test. When normality assumptions were not satisfied, non-parametric analysis was carried out using the Kruskal-Wallis test followed by Dunn’s post hoc test. If the data were normally distributed, group comparisons were made using one-way ANOVA with Tukey’s post hoc correction for multiple comparisons. Results are reported as mean ± standard deviation (SD), unless specified otherwise. In graphical representations, line and bar charts show mean ± SD, while box-and-whisker plots indicate the full data range (minimum to maximum). Statistical testing was conducted using either one-way ANOVA with Tukey’s post hoc or Kruskal-Wallis with Dunn’s post hoc, depending on data distribution. Statistical significance is indicated by asterisks: * p ≤ 0.05, ** p ≤ 0.01, *** p ≤ 0.001, **** p ≤ 0.0001.

## 2. Results and Discussion

### 2.1 Particle morphology of INU-MMT hybrids following spray drying

SEM was used to assess the morphology of spray-dried INU-MMT hybrids and their precursor components. Raw INU appeared as highly agglomerated, flake-like clusters (20–50 µm, **Fig. 1A**), consistent with its amorphous structure and self-association via hydrogen bonding. This porous morphology contributes to INU’s well-documented water retention capacity and dispersion behavior in aqueous environments [31]. MMT particles displayed irregular, amorphous aggregates exceeding 100 µm in size (**Fig. 1B**). This morphology is characteristic of stacked aluminosilicate platelets, where adsorption activity is localized within interlayer spaces and platelet edges [23, 26]. In contrast, the physical mixture of INU and MMT (**Fig. 1C**) showed distinct spherical INU particles coexisting with fragmented MMT clusters, indicating minimal interaction in the dry-mixed state. However, the spray-dried INU-MMT hybrid (**Fig. 1D**) highlighted a notable shift in particle morphology, forming uniformly aggregated microparticles (∼5–10 µm). The transformation suggests that spray drying facilitated a degree of structural integration between the two components. This is consistent with prior reports where spray drying reduced particle size and improved homogeneity in polysaccharide–clay systems [26].

**Fig 1.**
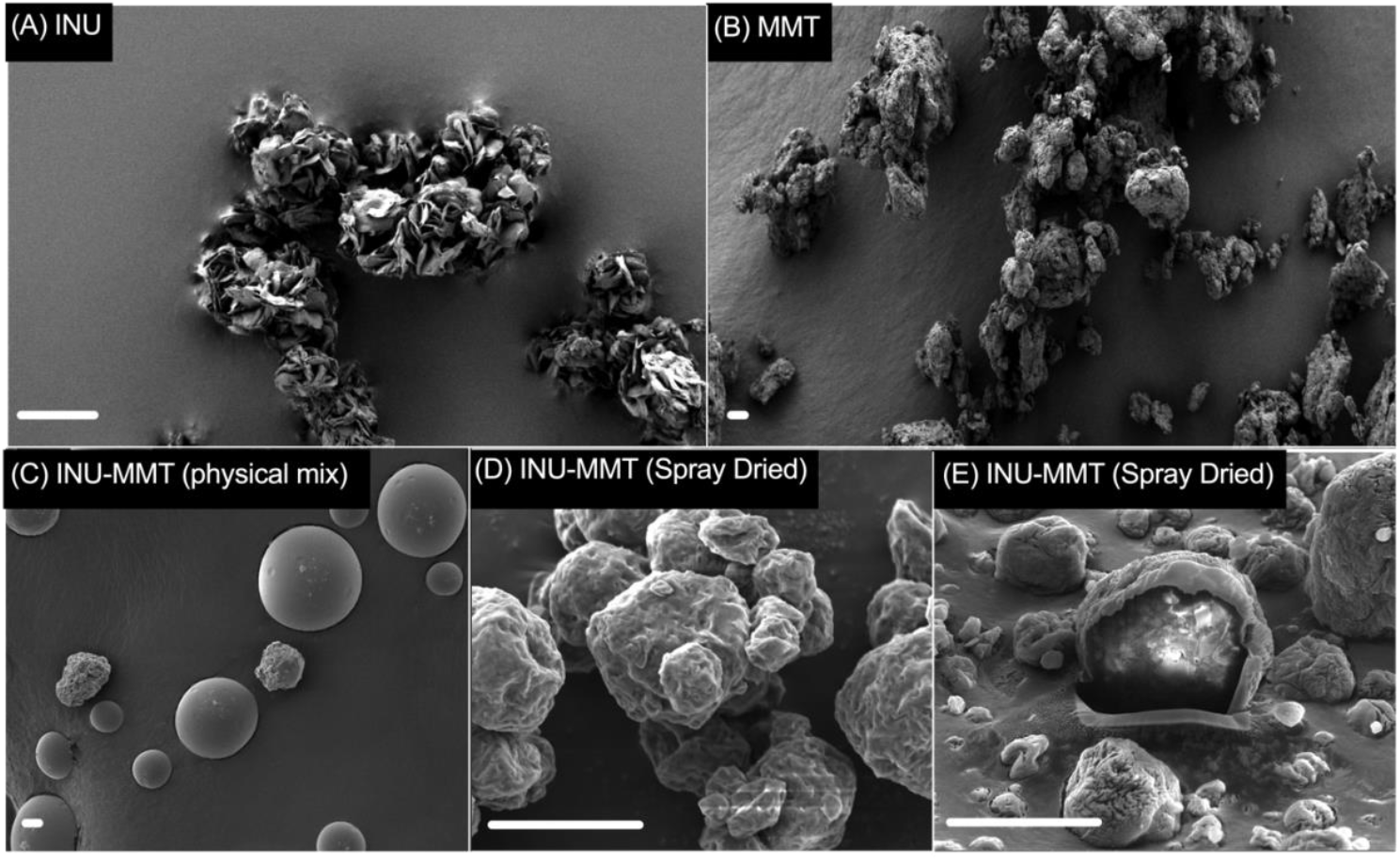
SEM images of INU-MMT hybrids and their precursors. (**A**) Raw INU appeared as amorphous, flake-like agglomerates (20–50 µm). (**B**) MMT demonstrated irregular, platelet-based aggregates (>100 µm). (**C**) The physical mixture revealed distinct INU and MMT particles with minimal interaction. (**D**) Spray-dried INU-MMT hybrids formed integrated microparticles (∼5–10 µm) with surface indentations. (**E**) Cross-sectional SEM (FIB-SEM) highlighted a partially porous internal structure. These features suggest spray drying enhanced particle integration and potential lipid-binding functionality. Scale bars = 10 µm.

Distinct surface indentations observed on the spray-dried hybrids (**Fig. 1D**) likely result from rapid solvent evaporation and polymer redistribution, a phenomenon commonly reported in spray-dried carbohydrate-based formulations [32]. As solutes accumulate at the droplet surface and form a rigid shell, internal capillary forces may lead to buckling, shaping the final particle morphology [33]. FIB-SEM cross-sectional imaging further revealed a partially porous internal architecture, which may enhance surface area and lipid-binding efficiency (**Fig. 1E**). These internal features are consistent with prior studies that associate porosity with improved functionality in spray-dried polysaccharide–clay hybrids [34].

Although the morphological changes suggest substantial structural integration, the current study did not include quantitative particle measurements such as size distribution, porosity index, or surface area. Future studies should incorporate techniques such as laser diffraction, BET surface analysis, to precisely characterize these particle features for optimizing structure–function relationships.

### 2.2. INU-MMT interaction with lipid droplets during lipolysis

To investigate the behavior of spray-dried INU-MMT hybrids during lipid digestion, confocal fluorescence microscopy was used to assess their interactions with lipid droplets during in vitro gastrointestinal lipolysis. INU-MMT hybrids were stained with rhodamine B (**Fig. 2A**, red), while medium-chain triglyceride (MCT) lipids were labeled with coumarin 6 (**Fig. 2B**, yellow). The overlay images revealed that after 60 minutes of digestion, INU-MMT particles clustered at the periphery of lipid droplets rather than fully dispersing within the lipid phase (**Fig. 2C**), suggesting surface-level adsorption rather than internal co-localization. This surface association is consistent with earlier findings that MMT limits lipid digestion by adsorbing FFAs through hydrogen bonding rather than intercalation within the clay interlayers [23]. These findings were supported by reduced interlayer spacing and disrupted platelet morphology observed post-lipolysis. The localization of INU-MMT around lipid droplets in this study mirrors these behaviors, suggesting adsorption may occur at broken platelet edges or external clay surfaces. Additionally, a similar study reported that MMT enhanced fecal lipid excretion by immobilizing dietary lipids *in vivo*, further substantiating its lipid-binding potential [25]. The mechanism likely involves electrostatic and hydrophobic interactions facilitated by the hybrid’s high surface porosity, as previously described with MMT [35]. In those systems, MMT selectively adsorbed FFAs and triglycerides, altering lipolytic product speciation and reducing their partitioning into solubilized phases, which could explain the reduced lipid bioavailability seen in similar formulations. Overall, the observed interactions support the hypothesis that spray-dried INU-MMT hybrids act as a lipid-binding interface during digestion. However, future studies are warranted to quantify lipid and FFA adsorption kinetics on the INU-MMT hybrid.

**Fig 2.**
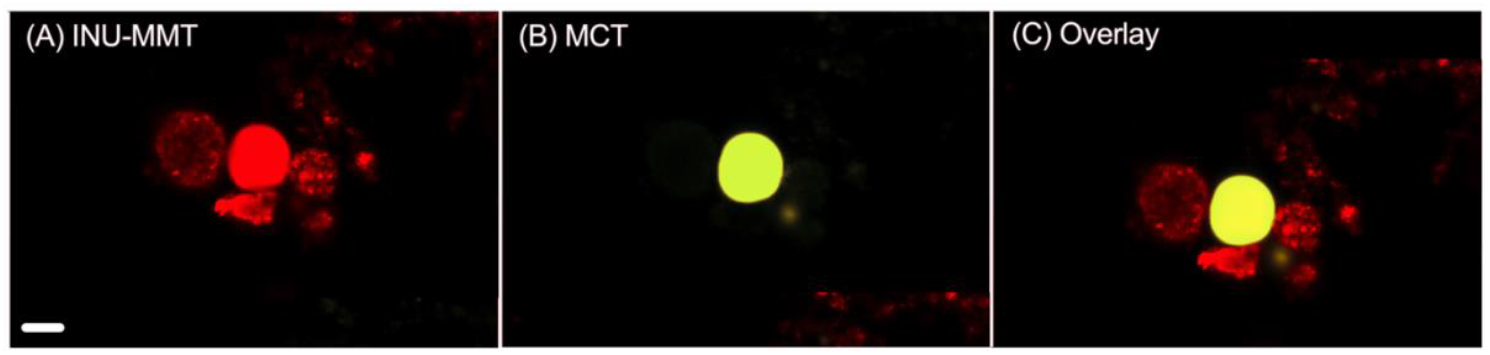
Confocal fluorescence microscopy images illustrating the interaction between rhodamine B-stained INU-MMT hybrids (**A**, red) and coumarin 6-stained MCT lipid droplets (**B**, yellow) following 60 minutes of in vitro lipolysis under biorelevant intestinal conditions. The merged image (**C**) reveals aggregation of INU-MMT around lipid droplets, suggesting surface-level adsorption of lipolytic products rather than internal co-localization. Scale bar = 10 µm.

### 2.3 Reduced FFA release by INU-MMT in simulated digestion conditions

The influence of INU-MMT on lipid digestion was evaluated under simulated fed-state intestinal conditions using MCT as a model dietary lipid. The MMT mechanisms of FFA release restriction has been postulated being via adsorbing lipolysis products or interfering with *lipase* enzymatic access at the lipid-in-water interface, thereby inhibiting hydrolysis of triglycerides into FFAs [36, 37]. After 60 minutes of *in vitro* lipolysis, untreated MCT released 51.33 ± 0.08 mmol/L of FFAs. This was significantly reduced in the presence of INU (35.05 ± 0.08 mmol/L; 32% reduction), MMT (17.29 ± 0.04 mmol/L; 66% reduction), and INU-MMT (18.27 ± 0.08 mmol/L; 64% reduction) (**Fig. 3B**). One-way ANOVA with Tukey’s post hoc test confirmed significant differences between all groups (p < 0.0001). While INU-MMT achieved a similar reduction in FFA release as MMT alone, it significantly outperformed INU (p < 0.0001). The preservation of MMT’s lipid-inhibitory function within the hybrid is particularly important, as INU is anticipated to support metabolic health through alternative mechanisms independent of lipid digestion. These complementary actions are postulated to provide additive therapeutic benefits. The kinetic profiles of FFA release (**Fig. 3A**) demonstrated that MCT alone rapidly reached a plateau by 40 min, consistent with complete lipolysis. INU exhibited a similar trend, albeit at a lower rate. In contrast, both MMT and INU-MMT exhibited early plateaus within 10–15 min, consistent with prior studies suggesting that MMT impedes lipase access to the lipid-water interface by adsorbing lipolytic products or calcium ions (Ca^2+^), which are essential for soap formation and FFA solubilization [23]. These mechanisms may restrict interfacial lipid digestion and contribute to the observed rapid suppression in FFA release [38]. This was reflected by a significant reduction in the digestion rate constant when digestion kinetics were fitted into a site-filling model, suggesting restricted lipid hydrolysis [39].

**Fig 3.**
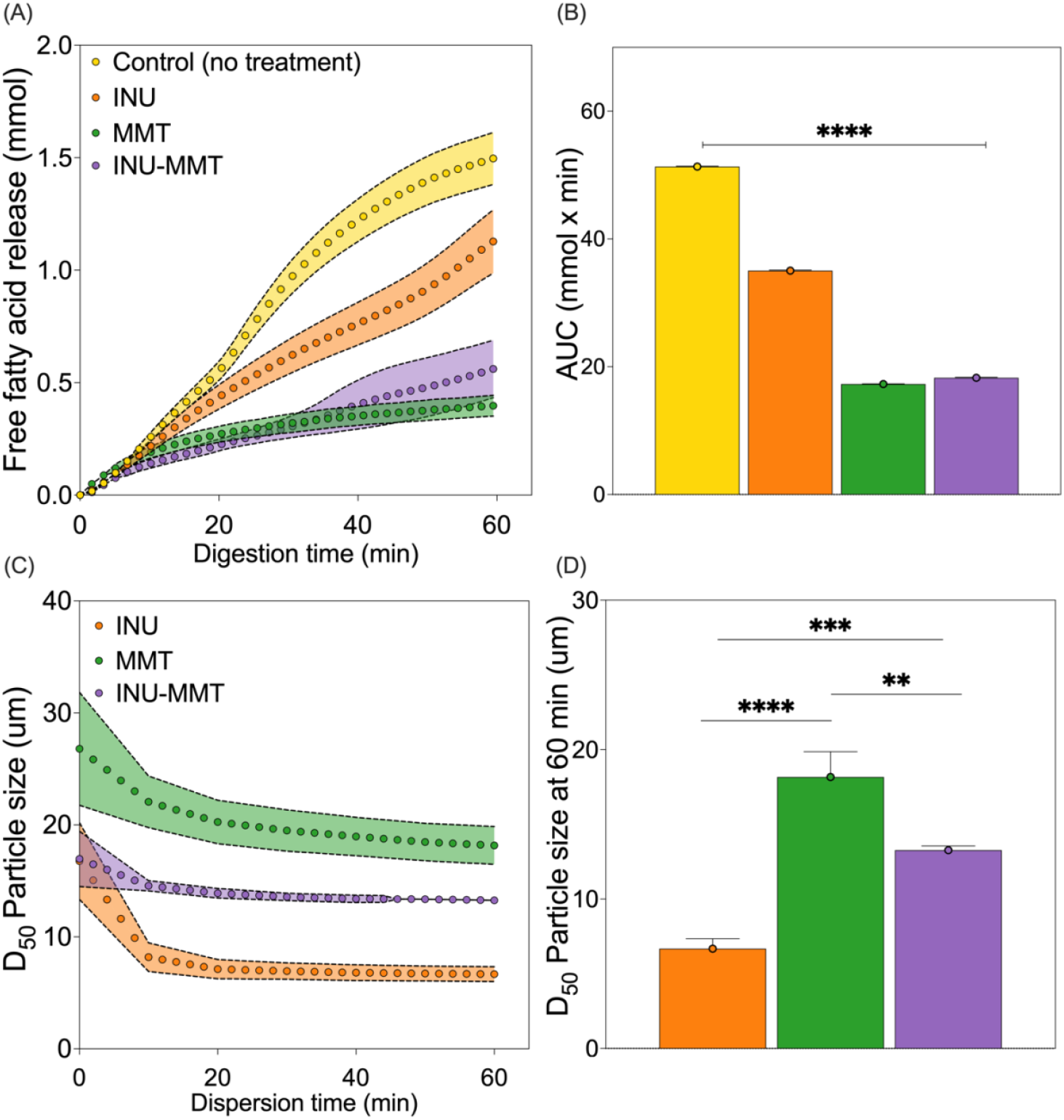
(**A**) INU-MMT and MMT sharply plateaued FFA release, indicating strong lipolysis inhibition. (**B**) AUC analysis showed 64–66% FFA reduction with INU-MMT and MMT, significantly greater than INU alone. (**C**) Particle dispersion in pH 6.5 buffer over 60 min showed stability differences across groups. (**D**) Final particle sizes revealed INU-MMT (13.27 ± 0.29 µm) was smaller than MMT (18.17 ± 1.69 µm) but larger than INU (6.68 ± 0.67 µm), suggesting moderate colloidal stabilization by the hybrid. Data are reported as mean ± SD, except for particle size data, reported as median (D_50_) ± SD (n = 3). Group differences were evaluated using one-way ANOVA followed by Tukey’s post hoc test. Statistical significance is denoted as follows: * p ≤ 0.05, ** p ≤ 0.01, *** p ≤ 0.001, **** p ≤ 0.000).

Redispersion behavior in small intestinal conditions (pH 6.5) was also assessed over 60 minutes (**Fig. 3C & 3D**). INU remained colloidally stable with a final particle size of 6.68 ± 0.67 µm, while MMT exhibited strong aggregation with significantly larger particles (18.17 ± 1.69 µm). INU-MMT particles exhibited an intermediate redispersion (13.27 ± 0.29 µm), which was significantly smaller than MMT alone (p = 0.0032), but not standalone INU (p = 0.0007). The low colloidal stability of MMT is likely due to its different charge characteristics at varying pH levels. At pH 6.5, the particles tend to aggregate due to charge interactions between the edges and faces of the clay layers, leading to larger particle sizes and reduced colloidal stability [40]. The current study highlights that hybridization with INU confers moderate colloidal stabilization to MMT via improved dispersion or reduced agglomeration. The improved redispersibility of INU-MMT may enhance its uniform distribution in intestinal fluid, promoting greater gut-activities for its individual components. The reduction in particle size over time for INU-MMT further implies partial disaggregation and dissolution of INU components [41]. These findings are consistent with earlier observations from SEM (**Fig. 1**), where spray-dried INU-MMT exhibited smaller, more uniform morphologies compared to MMT alone.

### 2.4. Restoration of HFD-induced gut microbiota composition by INU-MMT

To investigate whether INU-MMT modulates gut microbial communities altered by HFD, 16S rRNA gene sequencing was conducted on fecal samples collected at day 21. Analysis of α-diversity metrics indicated that INU-MMT increased microbial richness and evenness relative to HFD, albeit with varying statistical significance across indices (**Fig. 4A**). While Chao1 richness (67.95 ± 15.61 vs. 51.48 ± 11.78) exhibited a 32% increase, the difference was not statistically significant (p = 0.47), suggesting only a modest expansion in species number. Similarly, the Shannon index rose by 13% in the INU-MMT group (2.38 ± 0.34 vs. 2.11 ± 0.19), but significance was only reached when compared to INU alone (p = 0.041). Importantly, the Simpson index (0.76 ± 0.08 vs. 0.60 ± 0.08; p = 0.016) revealed a significantly more even distribution of microbial taxa in INU-MMT-treated rats compared to HFD, indicating a balanced microbiome with reduced dominance by specific groups. These findings align with studies demonstrating INU’s capacity to enhance α-diversity and counteract HFD-induced dysbiosis [7]. Notably, INU has been demonstrated to enrich *Bifidobacterium, Akkermansia*, and SCFA-producing *Ruminococcaceae*, all of which were elevated in the current INU-MMT group [42].

**Fig 4.**
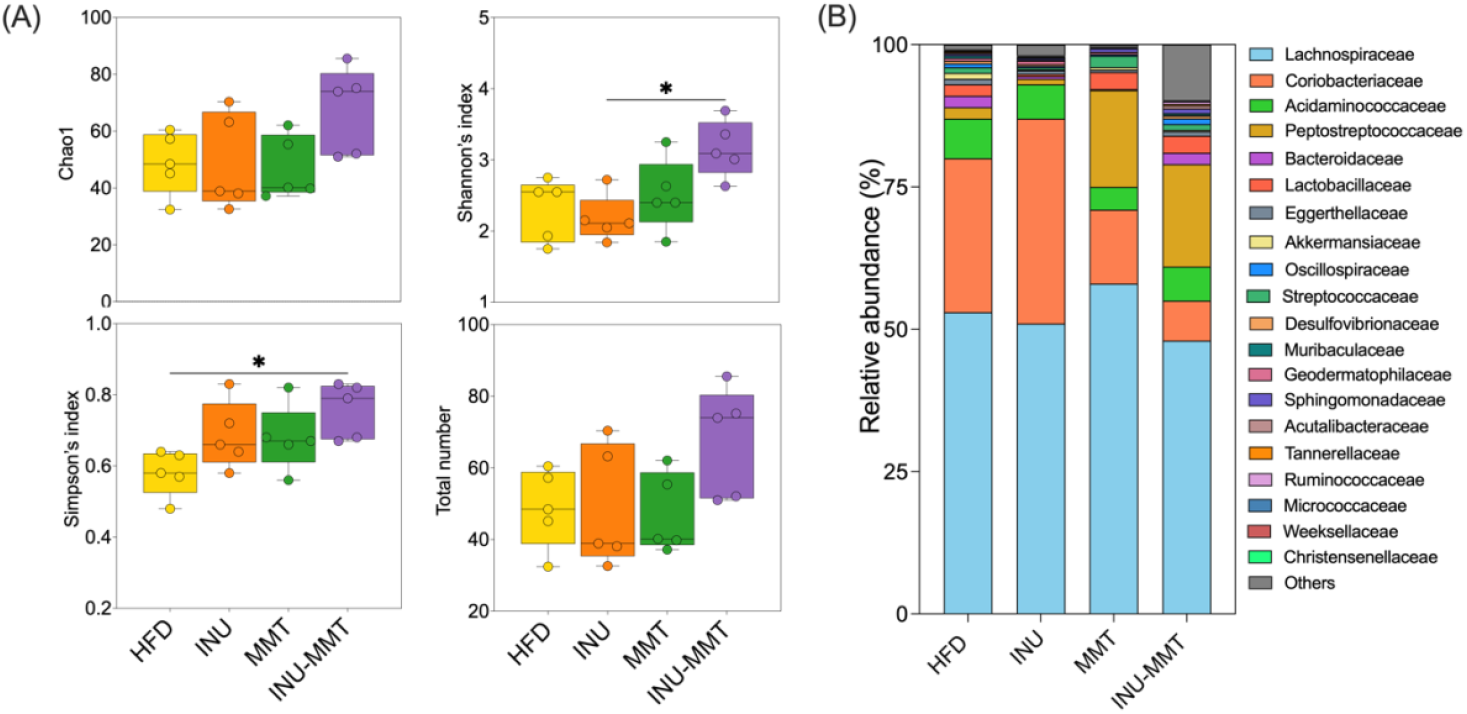
Impact of INU-MMT and precursor treatments on gut microbiota composition at day 21. 338 (**A**) INU-MMT significantly enhanced microbial diversity, as indicated by increased Simpson’s 339 index (p ≤ 0.05), while no significant differences were detected in Chao1, Shannon, or observed 340 species metrics. (**B**) At the family level, INU-MMT markedly enriched beneficial taxa 341 associated with SCFA production and mucosal health, including *Peptostreptococcaceae* (8-342 fold), *Ruminococcaceae* (3.5-fold), *Eggerthellaceae* (7.7-fold), and *Akkermansiaceae* (2.5-343 fold), compared to the HFD group. Data are presented as mean ± SD (n = 5 per group). 344 Statistical analysis was performed using Kruskal–Wallis test with Dunn’s multiple 345 comparisons. Statistical significance is denoted as follows: * p ≤ 0.05, ** p ≤ 0.01, *** p ≤ 346 0.001, **** p ≤ 0.0001. 347

At the taxonomic level, INU-MMT distinctly reshaped the gut microbiota composition (**Fig. 4B**). The HFD group was dominated by *Lachnospiraceae* (53% relative abundance), a family implicated in intestinal barrier dysfunction and metabolic endotoxemia through long-chain fatty acid production by members such as *Fusimonas intestini* [43, 44]. INU-MMT reduced *Lachnospiraceae* abundance by 9.4%, counteracting this pro-inflammatory profile. *Peptostreptococcaceae*, linked to fiber fermentation and butyrate generation, increased from 2% in HFD to 18% in INU-MMT, representing an 8-fold increase. Similarly, *Ruminococcaceae* abundance rose 3.5-fold, supporting mucosal integrity through SCFA production and immunomodulation [45, 46]. *Eggerthellaceae* and *Akkermansiaceae* - families often linked with polyphenol metabolism and mucin degradation, respectively - were also significantly enriched in the INU-MMT group, by 7.7-fold and 2.5-fold. These taxa are central to maintaining gut barrier function and host metabolic balance, and their increase likely reflects synergistic effects of the hybrid compound. Prior work has shown that INU promotes *Akkermansia muciniphila*, a hallmark of a balanced gut microbiome, with roles in glucose regulation and intestinal barrier enhancement [42]. Interestingly, while MMT alone increased *Peptostreptococcaceae* (to 17%) and *Eggerthellaceae* (to 0.443%), its effects were consistently less pronounced than those of the hybrid. These differences suggest that INU-MMT may possess unique structural properties that enhance colonization and activity of beneficial taxa, though this has not been studied yet.

### 2.5 MMT-based treatments alter overall microbial community structure distinct from its precursors

Beta diversity metrics were used to assess how overall microbial community composition varied between treatment groups. Unlike alpha diversity, which reflects diversity within individual samples, beta diversity compares differences between samples, highlighting the presence of distinct microbial communities. This was visualized using PCoA plots based on Bray–Curtis, Jaccard, and Euclidean distances (**Fig. 5**). A pseudo-F statistic measures variance between groups relative to within-group variance, with higher pseudo-F values indicating a greater separation between groups. Moreover, Bonferroni-adjusted p-values were applied to control for multiple comparisons and reduce the likelihood of false positives.

**Fig 5.**
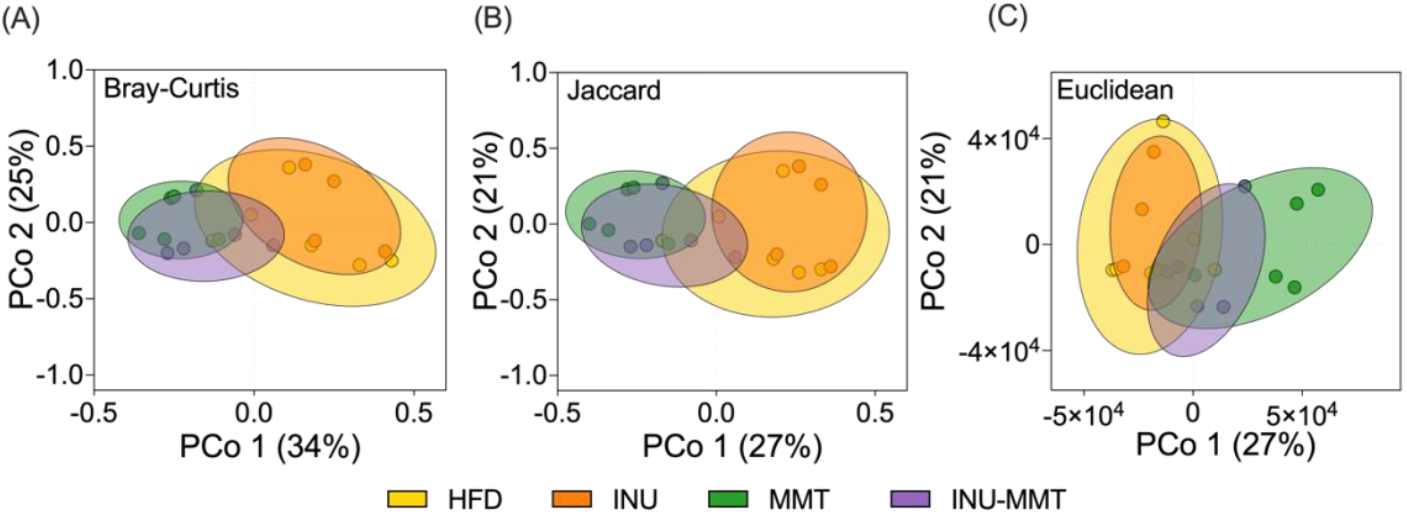
Principal coordinate analysis (PCoA) plots depicting beta diversity of gut microbiota at day 21 using (**A**) Bray–Curtis, (**B**) Jaccard, and (**C**) Euclidean distance metrics (n = 5 per group). INU-MMT differed significantly from INU across all metrics (adjusted p < 0.05), with Bray–Curtis and Euclidean distances also separating INU-MMT from HFD. MMT contributed most strongly to these shifts. Statistical differences were evaluated using PERMANOVA with pseudo-F values and Bonferroni-adjusted p-values. Ellipses represent 95% confidence intervals.

Based on Bray–Curtis distances, which measure both presence and abundance of taxa, INU-MMT was significantly different from the HFD group (pseudo-F = 2.98, p = 0.0108; adjusted p = 0.065), and from INU (pseudo-F = 5.16, p = 0.0022; Bonferroni-adjusted p = 0.013) (**Fig. 5A**). MMT also significantly differed from both HFD (pseudo-F = 4.60, adjusted p = 0.039) and INU (pseudo-F = 7.71, adjusted p = 0.013), confirming that the clay component has a strong effect on microbial community structure. Jaccard distances, which account for the presence or absence of taxa regardless of abundance, exhibited a divergent trend. INU-MMT differed significantly from INU (pseudo-F = 3.50, adjusted p = 0.026), but not HFD (pseudo-F = 2.18, adjusted p = 0.078). MMT again diverged significantly from both HFD and INU (adjusted p < 0.03) (**Fig. 5B**). Euclidean distance analysis, based on absolute differences in microbial abundances, resulted in similar findings to that of Jaccard (**Fig. 5C**). INU-MMT was again significantly different from INU (pseudo-F = 4.61, adjusted p = 0.039), but MMT resulted in a non-significant separation from both HFD and INU (pseudo-F > 7.4, adjusted p < 0.06). Notably, INU-MMT did not differ significantly from MMT across any metric, suggesting MMT exerts a dominant influence on the overall microbial profile of the hybrid. These results indicate that INU-MMT shifts the overall composition of the gut microbiota in a way that is distinct from INU and HFD, but largely driven by its clay component. Other studies have also confirmed MMT treatment in diet-induced obesity models significantly altered gut microbiota, decreasing *Firmicutes* and increasing *Bacteroidetes* (i.e., F/B ratio), changes that are positively correlated with metabolic health [24, 47]. Mice fed a 3% (w/w) MMT-supplemented diet for 14 days also demonstrated significant changes in beta diversity based on weighted UniFrac analysis [25]. These findings highlight that INU-MMT’s microbial modulation is not limited to changes in a few specific taxa, but rather alters a widespread selection of taxa.

### 2.6 INU-MMT suppresses pro-inflammatory and lipolytic microbial functions via reduced predicted enzyme abundance

PICRUSt2 was used to infer functional shifts in microbial enzymes from 16S rRNA gene profiles at day 21. A key finding was a marked reduction in predicted lipopolysaccharide (LPS) N-acetylglucosaminyltransferase abundance - a core enzyme in LPS biosynthesis. Compared to HFD, treatment with INU, MMT, and INU-MMT reduced its abundance by 93%, 62%, and 74%, respectively (**Fig. 6A**), although only the INU group reached statistical significance (p = 0.0007). LPS is a potent endotoxin derived from gram-negative bacteria that activates toll-like receptor 4 (TLR4), triggering metabolic endotoxemia and systemic low-grade inflammation— a key driver in obesity and insulin resistance [48, 49]

**Fig 6.**
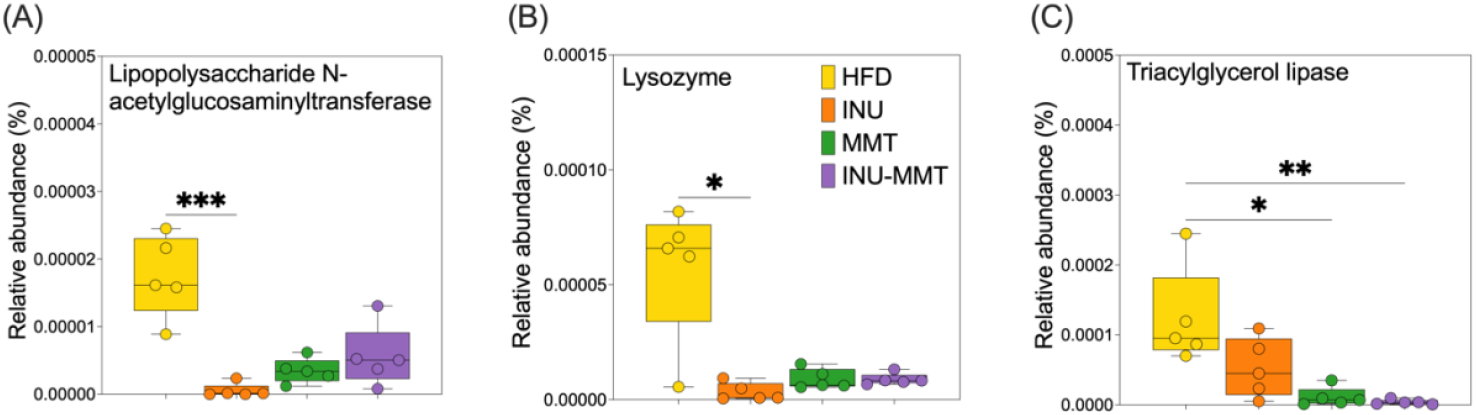
Predicted microbial enzyme abundance (PICRUSt2) in fecal samples (n = 5 per group). INU-MMT reduced levels of (**A**) LPS N-acetylglucosaminyltransferase (74%), (**B**) lysozyme (81%), and (**C**) significantly reduced triacylglycerol lipase (98%, p = 0.0067) compared to the HFD, indicating reduced pro-inflammatory and lipolytic potential. Statistical analysis was performed using Kruskal–Wallis test with Dunn’s multiple comparisons. Statistical significance is denoted as follows: * p ≤ 0.05, ** p ≤ 0.01, *** p ≤ 0.001, **** p ≤ 0.0001.

The reduction in this LPS-linked enzyme across all treatments suggests a microbial shift away from gram-negative taxa. This is consistent with the microbial properties of INU, which selectively enriches beneficial genera such as *Akkermansia*, which is linked to suppressed LPS biosynthesis and improved metabolic profiles [50]. Another important enzyme suppressed by all treatments was lysozyme, an antimicrobial protein often elevated during microbial turnover and adipose inflammation. INU, MMT, and INU-MMT treatments resulted in reductions of 87%, 84%, and 81%, respectively (**Fig. 6B**), with only INU achieving statistical significance (p = 0.0139). Elevated lysozyme levels in adipose tissue have been linked to impaired adipogenesis and chronic inflammation in HFD models [51]. Moreover, triacylglycerol lipase abundance, responsible for microbial lipid hydrolysis, was profoundly reduced by INU-MMT (98%), MMT (90%), and INU (67%) (**Fig. 6C**). Significant suppression was observed in both MMT (p = 0.0385) and INU-MMT (p = 0.0067) compared to HFD. This aligns with prior studies showing that MMT can adsorb dietary lipids, thus physically limiting lipid accessibility to microbial and host lipases [23, 25, 26, 37]. INU-MMT may enhance this effect by limiting MMT aggregation in pH 6.5 media (**Fig. 3D**), yielding smaller, more dispersed particles with increased interfacial interaction with lipolytic products.

### 2.7. INU-MMT attenuates weight gain in HFD-induced obese rats

Daily treatment with INU-MMT over 21 days markedly reduced weight gain in high-fat diet (HFD)-fed rats (**Fig. 7A**). Cumulative weight gain, assessed via AUC analysis, was 4.7% lower in the INU-MMT group compared to HFD alone, which was significantly greater than the modest reductions achieved by INU (2.0%) and MMT (1.5%) individually (**Fig. 7B**). Statistical comparison using Tukey’s post hoc test confirmed the superiority of INU-MMT over all other groups (p < 0.0001 vs HFD, INU, and MMT). This attenuation in weight gain by INU-MMT likely reflects gut-activity via alternate mechanisms. These mechanisms may include restricted lipid digestion, as well as eubiotic shifts within gut microbiota. In an earlier study, diet-induced obese rats supplemented with MMT (1g/kg bodyweight/d) were found to have significantly attenuated weight gain of 1.9% [22]. Similarly, another study reported that diet-induced obese rats exhibited up to a 3.8% reduction in weight gain across 21 days of daily treatment with INU [7]. Together, the synergy of the INU-MMT hybrid may contribute to slower weight accumulation in HFD environments.

**Fig 7.**
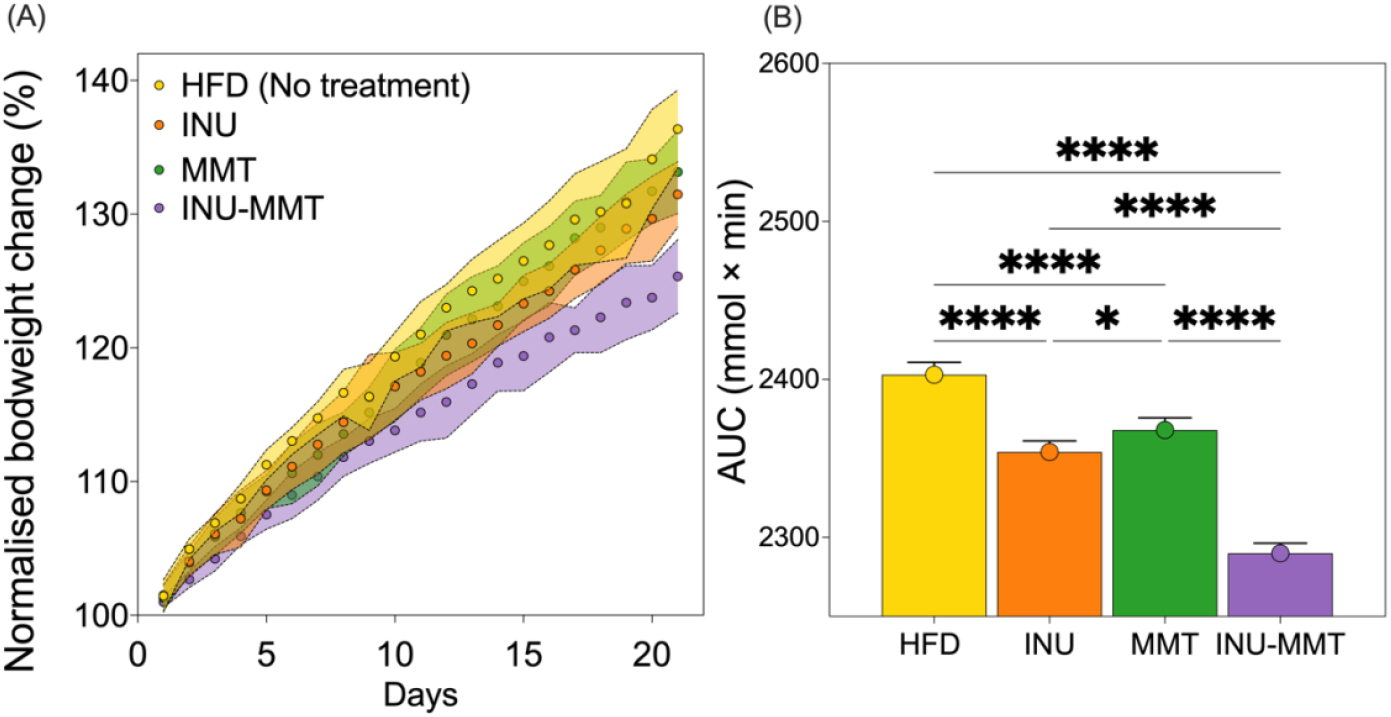
Body weight gain in HFD-fed rats over 21 days of daily treatment (n = 5 per group). (**A**) INU-MMT-treated rats resulted in the greatest attenuation of weight gain compared to other treatment groups. (**B**) Area under the curve (AUC) analysis revealed that INU-MMT reduced cumulative weight gain by 4.7% relative to HFD, greater than INU (2.0%) or MMT (1.5%) alone. Statistical analysis was performed using one-way ANOVA followed by Tukey’s post hoc test for multiple comparisons. Significance was indicated as follows: * p ≤ 0.05, ** p ≤ 0.01, *** p ≤ 0.001, **** p ≤ 0.0001.

## Conclusion

Spray-dried INU-MMT hybrids effectively counteract the metabolic and inflammatory effects of high fat diet-induced obesity in rats. The INU-MMT hybrid reduced *in vitro* lipid digestion, improved *in vivo* gut microbial diversity, and shifted microbial composition toward taxa associated with short-chain fatty acid production and mucosal health. The hybrid also attenuated predicted pro-inflammatory and lipolytic microbial functions, indicating a broad impact on both gut microbial composition and function. Compared to INU or MMT alone, INU-MMT exerted synergistic effects in reducing weight gain in rats. These findings highlight the therapeutic potential of INU-MMT hybrids as multifunctional therapies or adjuvants for weight management and metabolic diseases. Future studies necessitate exploring the long-term efficacy of INU-MMT, and the potential to incorporate additional bioactives for enhanced clinical applications.

## Conflicts of interest

The authors declare no conflict of interest.

## Acknowledgments

This research was funded by the Australian Government through the Australian Research Council (LP230100345). P.J. acknowledges the Hospital Research Foundation (THRF) group (2022-CF-EMCR-004-25314) for their generous support of this project. The authors also acknowledge the facilities, scientific, and technical support provided by Microscopy Australia at the University of South Australia and Adelaide Microscopy at the University of Adelaide.

## Author contributions: CRediT

**Amin Ariaee:** Conceptualization, Methodology, Investigation, Formal analysis, Data curation, Visualization, Writing – Original draft preparation. **Alex Hunter:** Methodology, Investigation, Visualization, Writing – Reviewing and Editing. **Anthony Wignall:** Supervision, Writing – Reviewing and Editing. **Kristen Bremmell:** Writing – Reviewing and Editing. **Clive**

**Prestidge:** Supervision, Writing – Reviewing and Editing. **Paul Joyce:** Conceptualization, Funding acquisition, Supervision, Project administration, Writing – Reviewing and Editing.

## Data Availability Statement

The data for all findings of the current study is available at request from the corresponding author.

## Notes

### Competing Interest Statement

The authors have declared no competing interest.

